# Common lizards break Dollo’s law of irreversibility: genome-wide phylogenomics support a single origin of viviparity and re-evolution of oviparity

**DOI:** 10.1101/225086

**Authors:** Hans Recknagel, Nick Kamenos, Kathryn R. Elmer

**Affiliations:** Institute of Biodiversity, Animal Health & Comparative Medicine, College of Medical, Veterinary & Life Sciences, University of Glasgow, Glasgow, G12 8QQ UK; School of Geographical and Earth Sciences, University of Glasgow, Glasgow G12 8QQ, UK

**Keywords:** Squamata, Lacertidae, Dollo’s law, viviparity, biogeography, molecular systematics

## Abstract

Dollo’s law of irreversibility states that once a complex trait has been lost in evolution, it cannot be regained. It is thought that complex epistatic interactions and developmental constraints impede the re-emergence of such a trait. Oviparous reproduction (egg-laying) requires the formation of an eggshell and represents an example of such a complex trait. In reptiles, viviparity (live-bearing) has evolved repeatedly but it is highly disputed if oviparity has re-evolved. Here, using up to 194,358 SNP loci and 1,334,760 bp of sequence, we reconstruct the phylogeny of viviparous and oviparous lineages of common lizards and infer the evolutionary history of parity modes. Our phylogeny strongly supports six main common lizard lineages that have been previously identified. We find very high statistical support for a topological arrangement that suggests a reversal to oviparity from viviparity. Our topology is consistent with highly differentiated chromosomal configurations between lineages, but disagrees with previous phylogenetic studies in some nodes. While we find high support for a reversal to oviparity, more genomic and developmental data are needed to robustly test this and assess the mechanism by which a reversal might have occurred.

## 1. Introduction

There are numerous examples for the loss of a complex trait in the animal kingdom throughout evolution. Dollo’s law of irreversibility states that once such a complex trait has been lost, it cannot be regained (Dollo, 1893). Some exceptions to this rule have been discovered, though it remains a very rare phenomenon in evolution (Collin and Miglietta, 2008; Lynch and Wagner, 2010). Oviparity (egg-laying) is an example for such a complex trait and has been lost on several independent occasions throughout animal evolution (Lee and Shine, 1998; Murphy and Thompson, 2011). While there are more than a hundred independent transitions from oviparity to viviparity (live-bearing) in reptiles (Blackburn, 2006; Sites et al., 2011), only one robust example for the re-evolution of the eggshell is known to date (Lynch and Wagner 2010). Molecular mechanisms by which reversals in complex traits such as reproductive mode occur are to date unknown.

The common lizard (*Zootoca vivipara*) is the most widespread extant terrestrial reptile species. Its distribution covers nearly the whole of Europe, northern and central Asia and as far as Japan in its easternmost range. Within this distribution, common lizards have adapted to various extreme environments. Arguably the most salient of these adaptations is the evolution of viviparous, unique within the family of ‘true’ (lacertid) lizards that are otherwise oviparous. As one of the youngest transitions from oviparity (egg-laying) to viviparity (live-bearing) known in vertebrates (Pyron and Burbrink, 2014; Surget-Groba et al., 2006), common lizards are an emerging model system for the study of viviparity (Freire et al., 2003; Le Galliard et al., 2003; Murphy and Thompson, 2011). However, not all common lizards are live-bearing: of the six currently recognized common lizard lineages, two are oviparous and four are viviparous (Surget-Groba et al., 2006; Fig. 1). One oviparous lineage is restricted to northern Spain and southwestern France, allopatric to all other common lizard lineages. A second oviparous lineage occurs in the southern part of the Alps. Four viviparous lineages cover the rest of the Eurasian distribution (Mayer et al., 2000; Surget-Groba et al., 2006; Fig. 2).

**Figure 1.**
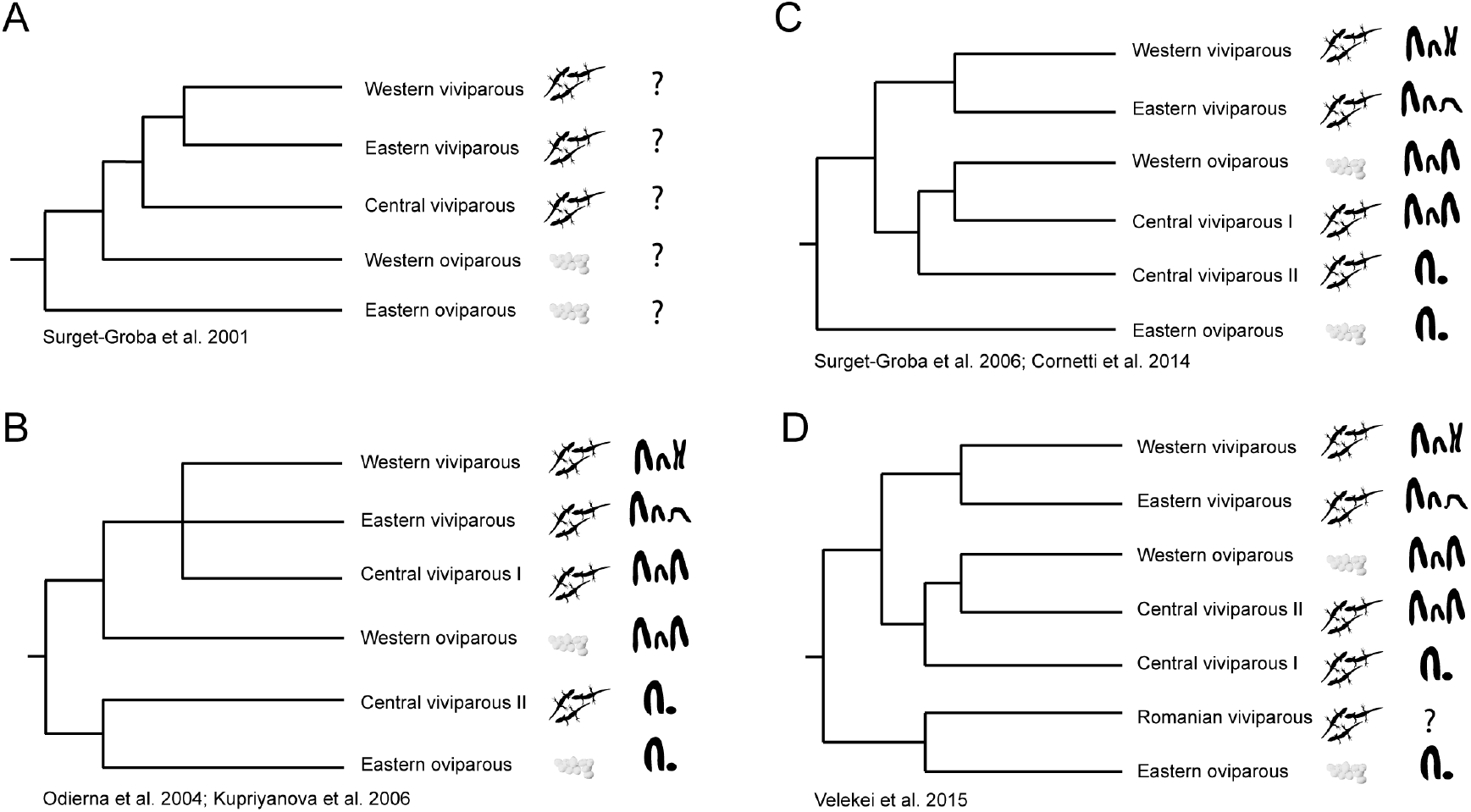
Alternative hypotheses for phylogenetic relationships of common lizards and parity mode evolution. Parity mode and sex chromosome configuration (ZW or Z_1_Z_2_W; Odierna et al., 2004) are illustrated next to each respective lineage. Phylogenetic tree A) involves a single origin of viviparity and was supported by one mtDNA gene. The second tree B) is based on karyological studies and suggests two independent origins of viviparity. Hypothesis C) suggests a reversal to oviparity as most parsimonious scenario, based on mtDNA and a few nuclear genes. The last phylogeny D) includes a recently discovered viviparous lineage in the Carpathians, which was found to be closely related to the most basal oviparous lineage. Parity mode evolution in this scenario involves two independent origins of viviparity and a reversal to oviparity.

**Figure 2.**
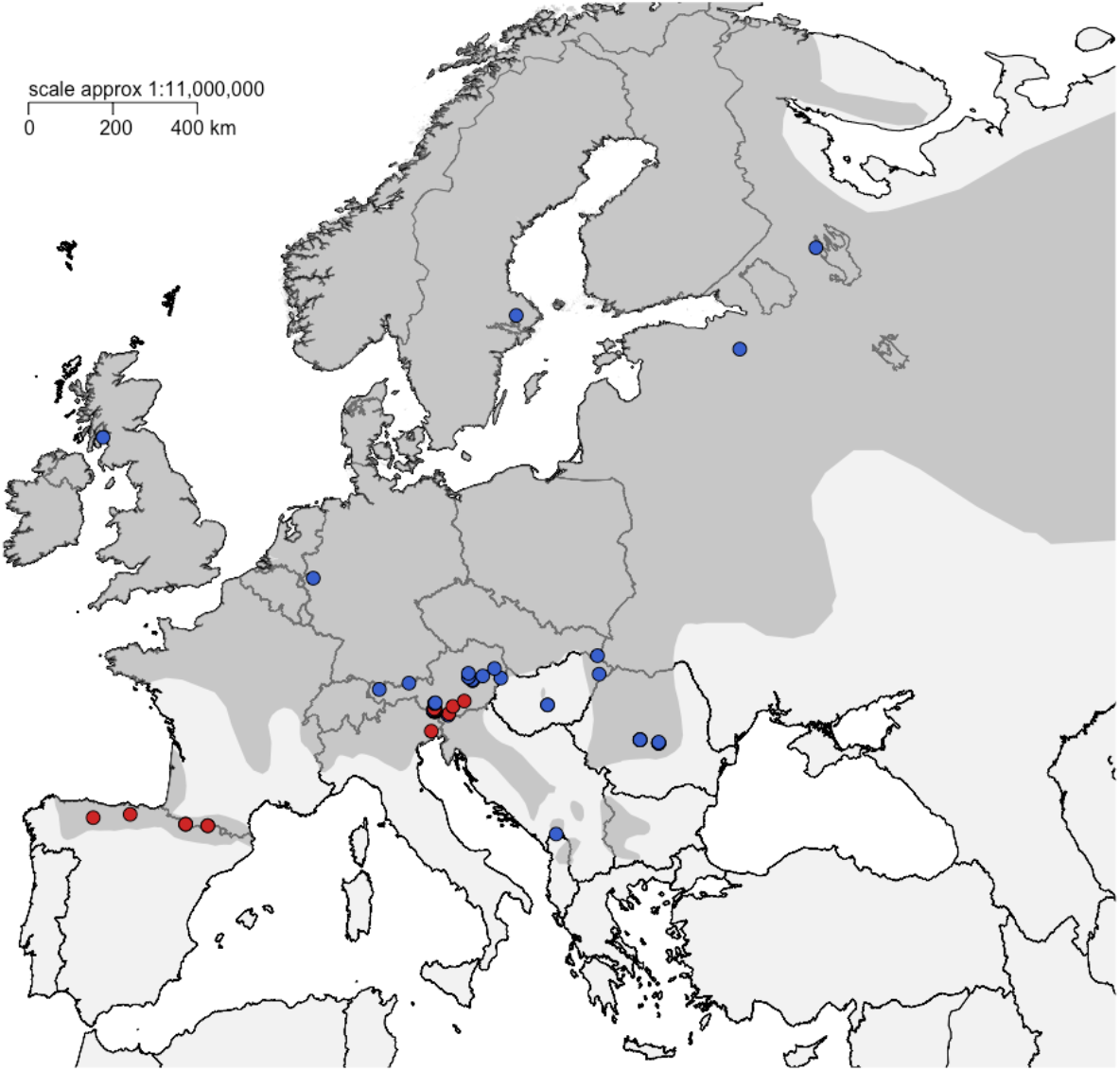
Map of common lizard (*Zootoca vivipara*) sampling locations within Europe. The dark grey shaded area marks the distribution of the common lizard in Europe. Each dot represents a single individual (red = oviparous; blue = viviparous) captured at the respective location. Note that a single individual from central Russia included in the phylogenetic analyses is outside the scope of the map (see Table S1).

The phylogenetic relationships within *Zootoca* have not been fully resolved. The evolutionary history of the two different parity modes has been controversial depending on which data was used to interpret the phylogenetic relationships. In a first study using a single mitochondrial gene, both oviparous lineages were found to be basal to all other viviparous lineages, consistent with a single origin of viviparity (Surget-Groba et al., 2001; Fig. 1A). However, subsequent analyses on the karyotype of common lizards resulted in a more complex evolutionary scenario, arguing for two origins of viviparity based on sex-chromosome evolution (Z_1_Z_2_W or ZW) (Odierna et al., 2004; Surget-Groba et al., 2006; Fig. 1B). More extensive geographic sampling and sequencing of mitochondrial genes instead favored a scenario of a single origin of viviparity followed by a reversal to oviparity in the Spanish Western Oviparous lineage (Cornetti et al., 2014; Surget-Groba et al., 2006; Fig. 1C), though this phylogeny was incompatible with a single origin of the Z_1_Z_2_W sex chromosome system. Finally, a population inhabiting the Carpathian region in Romania was discovered recently and was found to be most closely related to the phylogenetically basal Eastern Oviparous lineage based on mtDNA (Velekei et al., 2015; Fig. 1D). The reproductive mode of this lineage was not reported, but since all other common lizard populations in its geographic proximity are viviparous (Surget-Groba et al., 2006),this would suggest another independent origin of viviparity. However, all phylogenies to date have had limited support at basal nodes essential for the interpreting the evolutionary scenarios of parity mode evolution. Moreover, phylogenies reconstructed only from mitochondrial DNA have limited information and frequently misrepresent the ‘true’ phylogenetic relationships (Ballard and Whitlock, 2004; Near and Keck, 2013; Wallis et al., 2017). Therefore, it is essential to incorporate high resolution nuclear DNA sequencing to resolve difficult topologies. Moreover, coalescent-based approaches for disentangling incomplete lineage sorting effects and hybridization have considerably advanced phylogenetic reconstruction (Bouckaert et al., 2014; Pickrell and Pritchard, 2012; Posada, 2016).

The evolutionary implications for models involving several origins of viviparity and/or a reversal to oviparity are significant. A reversal to oviparity from viviparity is considered a very unlikely evolutionary scenario, presumably breaking Dollo’s law of irreversibility. Common lizard parity mode evolution could represent one of the very few examples for an exception to this rule (Surget-Groba et al., 2006). Further, the evolution of both oviparity and viviparity are difficult to study from a molecular genetic perspective because they have most frequently occurred at deep evolutionary time scales. Common lizards provide an example of recent parity mode changes and therefore a critical insight to usually more ancient evolutionary events.

To tackle this outstanding phylogenetic question, we use genome-wide phylogenomics with data from double-digest restriction-site associated DNA sequencing (ddRADSeq), a next generation sequencing (NGS) technique, to identify DNA polymorphisms across all common lizard lineages (Peterson et al., 2012; Recknagel et al., 2015, 2013). Using broad geographic sampling of 70 individuals, we reconstructed a nuclear phylogeny of 194,358 bp, and a mtDNA phylogeny based on cytochrome b, using coalescent, Maximum Likelihood, and Maximum Parsimony methods. We performed topological tests to assess likelihoods of alternative evolutionary scenarios for parity mode evolution based on our phylogenomic dataset, which consistently supported an evolutionary scenario. Our results strongly support a single origin of viviparity in common lizards and a subsequent reversal to oviparity in one derived lineage as the most parsimonious scenario of reproductive mode evolution.

## 2. Material and Methods

### 2.1 Sampling

Samples and specimens were obtained from the Natural History Museum in Vienna, the Royal Ontario Museum, and fieldwork during 2013-2016 (see Table S1 for specimens and Fig. 2 for a map of collecting localities). Lizards were collected by diurnal opportunistic searches. Tail clips (up to 2 cm) were extracted and preserved in 95-99% ethanol and lizards were released thereafter. Mode of reproduction was assessed by observation of an individual retained in captivity until oviposition/parturition or from data on other individuals at the same site.

### 2.2 Generation of molecular data

DNA was extracted from tissue using a Dneasy Blood and Tissue Kit (Qiagen) following the manufacturer’s protocol. Three genomic libraries were constructed using double-digest restriction-site associated DNA sequencing (ddRADSeq). The first two libraries were run on an IonProton sequencing machine with a median of 96 bp read length (ddRADSeq-ion; Recknagel et al., 2015) and the third library was paired-end sequenced on an Illumina HiSeq 4000 with 150 bp read length. Briefly, 1 ug of starting DNA material was digested using restriction enzymes *PstI-HF* and *MspI* and subsequently cleaned with the Enzyme Reaction Cleanup kit (Qiagen). Following purification, the amount of DNA in each individual was normalized to the sample with the lowest concentration within a library (237 ng in first, 400 ng in second, and 275 ng in third library) to minimize coverage variation. Platform specific barcoded (for IonProton: A-adapter, for Illumina: P1 adapter; binding to PstI-HF overhang) and global (for IonProton: P1-adapter, for Illumina: P2 adapter; binding to *MspI* overhang) adapters were ligated to the sticky ends generated by restriction enzymes. The ligated DNA fragments were then multiplexed and size-selected using a Pippin Prep (Sage Science) for a range between 175 – 225 bp for the IonProton platform and 150 – 210 bp for Illumina. To assure that the same set of loci are selected between platforms, size selection ranges were adjusted because adapter lengths are not the same between platforms. Seven separate PCR reactions (for details see Recknagel et al., 2015) were performed per genomic library and combined (Peterson et al., 2012). Following PCR purification, libraries were electrophoresed on a 1.25% agarose gel to remove any remaining adapter dimers and fragments outside the size range selected by the Pippin Prep. SYBRSafe (Life Technologies) was used for gel staining and bands in the size selected range were cut out manually and DNA was extracted from the matrix using a MinElute Gel Extraction Kit (Qiagen). Following the gel extraction, DNA was quantified using a Qubit Fluorometer with the dsDNA BR Assay. Quality and quantity of genomic libraries was assessed using a TapeStation or Bioanalyzer (Agilent Technologies). The first two libraries were sequenced at Glasgow Polyomics using an Ion PI Sequencing 200 Kit v3 on an Ion Proton PI chip at a target read size of 100 bp. The third library was sequenced at Edinburgh Genomics on an Illumina HiSeq 4000 machine with paired-end sequencing of 150 bp reads.

In addition to ddRADseq, mitochondrial DNA (mtDNA) from cytochrome b with primers MVZ04H and MVZ05L (~430 bp) was amplified (Smith and Patton, 1991) and PCR products were sequenced with the forward primer (MVZ04H) on an ABI 3130x at Dundee University. Sequences were quality checked by eye, and trimmed and aligned using Geneious v. 7.1.9 (Kearse et al., 2012). Data are deposited in NCBI (Genbank accession with manuscript acceptance).

### 2.3 Bioinformatic analysis

All NGS generated reads were analyzed using the RADseq software tool STACKS v.1.41 (Catchen et al., 2011). Reads were trimmed to a common length of 70 bp to maximize the number and length of retained reads (Recknagel et al., 2015). Libraries were de-multiplexed and all reads were sorted into stacks of loci within each individual (maximum distance of 2 bp within a locus). The minimum coverage threshold per individual locus was set to five. Each individual was then aligned to a *Zootoca vivipara* reference genome v. 0.9 (Yurchenko et al. in prep) using bwa (Li and Durbin, 2010) and samtools (Li et al., 2009). A catalogue of all loci identified across individuals was subsequently created using the genome referenced stacks from each individual.

Missing data can have a substantial impact on phylogenetic inference from NGS generated data and can vary between taxonomic and phylogenetic levels (Eaton et al., 2017; Jiang et al., 2014; Rowe et al., 2011; Streicher et al., 2016). Therefore, it is crucial to first evaluate the impact of missing data before phylogenetic analysis. We filtered our data with two main options: i) using a variable minimum number of individuals that a locus had to be present in, and ii) varying the number of SNPs per locus from one to three. The amount of missing data was increased from 0% to 90% at 10% intervals. For each of these categories, loci containing only a single SNP, two SNPs, three SNPs and one to three SNPs were extracted from the whole dataset. These datasets were extracted to test the impact of missing data and number of SNPs on phylogenetic resolution and to assess optimal settings for data extraction.

### 2.4 Phylogenetic analysis

Suitability of data sets that differed in degree of missing data and number and type of SNP loci was assessed by comparing the sum of bootstrap supports (at deep, at shallow, and at all nodes combined) (Huang and Lacey Knowles, 2016). The best performing dataset for inferring the evolutionary history of parity mode in common lizards was identified and chosen for more exhaustive phylogenetic and comparative analyses. This best performing dataset was assessed by constructing Maximum-likelihood (ML) phylogenies using the software RAxML vers. 8.1.20 with a GTRGAMMA substitution model of evolution (Stamatakis, 2006). Conditions producing the highest bootstrap sum phylogeny were the ones chosen for all subsequent analyses.

We inferred Maximum-likelihood (ML) phylogenies using RAxML. An initial phylogenetic analysis including the outgroup species *Iberolacerta horvathi* identified the Eastern Oviparous clade as basal to all five other *Zootoca* lineages with high confidence (bootstrap support 100), as has been shown by previous analyses (Cornetti et al., 2014; Mayer et al., 2000; Surget-Groba et al., 2006). We further used ADMIXTURE (vers. 1.3.0; Alexander et al., 2009) to test for monophyly of the main *Zootoca* lineages. ADMIXTURE assesses the genomic ancestry of individuals according to a given set of genetic clusters. A variable number of genetic clusters *k* was run, from 1 to 6 *k* and best fit inferred from ten-fold cross-validation. The genetic cluster with the lowest cross-validation error was chosen as optimal *k*. These analyses confirmed monophyly of the six main lineages and limited levels of admixture. Pairwise genetic differentiation between lineages was assessed using the R package diveRsity (Keenan et al., 2013).

A Maximum likelihood bootstrap search with 100 replicates using a GTRGAMMA model was performed in RAxML. Support values were drawn on the best scoring ML tree. The best ML tree was compared to four alternative pre-defined topologies, which had been proposed in previous studies. These topologies included i) both oviparous lineages basal to all viviparous lineages (Mayer et al., 2000; Surget-Groba et al., 2001; Fig. 1A) ii) Eastern oviparous lineage basal + Central viviparous II basal to all remaining viviparous and oviparous (Odierna et al., 2004; Surget-Groba et al., 2006; Fig 1C), iii) Eastern oviparous lineage basal + Central viviparous I basal to all remaining viviparous and oviparous lineages, and iv) Romanian lineage sister to Eastern oviparous and basal to all other lineages (Velekei et al., 2015). We computed per site log likelihoods for each of the five trees and used these to perform Approximately Unbiased tests (AU tests) (Shimodaira, 2002), Shimodaira-Hasegawa tests (SH tests) (Shimodaira and Hasegawa, 1999), Kishino-Hasegawa tests (KH tests), and Bayesian posterior probabilities (PPs) calculated by the BIC approximation as all implemented in CONSEL vs. 0.1a (Shimodaira and Hasegawa, 2001).

We performed a coalescent-based Bayesian approach to infer the topology in BEAST2 (Bouckaert et al., 2014). For this approach, we included a full alignment of all RAD loci (19,068 RAD loci; 1,334,760 total bp; 84,017 variant sites). The number of total SNPs differs from other analyses as loci were set to be present in at least 40% of individuals of each of the six lineages, instead of just being present in at least 40% of individuals across the whole phylogeny. We used the GTRGAMMA substitution model. The analysis was run on CIPRES (Miller et al., 2010) for 500 million generations sampling trees every 50,000 and discarded 10% as burn-in. Convergence was assessed in TRACER (Rambaut and Drummond, 2009) and accepted if ESS values of all parameters were larger than 100.

Additional phylogenetic analyses were carried out under the Maximum Parsimony (MP) optimality criteria. We performed a heuristic bootstrap search with 2000 replicates carried out in PAUP* (Swofford, 2002) using TBR branch swapping and with ten random addition sequence replicates for each bootstrap replicate. The 50% consensus bootstrap tree was compared to phylogenies generated with ML and Bayesian analyses.

To incorporate potential past migration events and incomplete lineage sorting effects, we performed a TREEMIX v.1.3 (Pickrell and Pritchard, 2012) search using only independent SNPs (one SNP per locus; 49,107 loci included) and a window size of 1000 bp. We included zero to six migration events and compared the variance explained between resulting tree with and without migration events to evaluate the impact of migration. We calculated f3-statistics to assess whether admixture has played a role in the evolution of common lizard lineages.

For the mitochondrial dataset, we performed a bootstrap ML search using RAxML (100 bootstrap replicates), MP using the same parameters mentioned above and Bayesian reconstruction with BEAST2 to generate the phylogeny. The best substitution model for BEAST2 was inferred from eleven different substitution schemes in JMODELTEST2 (Darriba et al., 2012) based on lowest AICc and run on CIPRES. We ran BEAST2 for 20 million generations and discarded 10% as burn-in. Convergence was inferred if ESS values in TRACER were larger than 100.

## 3. Results

### 3.1 Data evaluation and identification of optimal parameters for phylogenomic dataset

Total number of generated reads was 828,000,972 (1^st^ library: 10,000,000 reads, 2^nd^ library: 42,377,658 reads, 3^rd^ library: 775,623,314 paired-end reads). After sorting reads into individual loci, mean coverage per individual was 27.6x with a standard deviation of 11.0x (range: 9.2x – 66.9x; median: 24.1x).

We found that phylogenetic resolution generally improved by accepting larger amounts of individuals with missing data (Fig. S1). The best summed bootstrap support was achieved using loci that were present in at least 40% of all individuals. Accepting more missing data this did not improve phylogenetic resolution. The highest number of SNPs (including up to three SNPs) resulted in the overall highest phylogenetic resolution (Fig. S1). Therefore, we chose the dataset with loci present in at least 40% of all individuals and including all SNPs (no restriction on number of SNPs per locus) for all subsequent analyses. Genotyping error was low (2.0-2.9% per SNP) based on three technical replicates and comparable to previous studies (Mastretta-Yanes et al., 2015; Recknagel et al., 2015).

### 3.2 Mitochondrial DNA phylogeny

The final alignment of the cytochrome b gene consisted of 428 bp (42 parsimony informative sites). HKY+I was identified as the best substitution model for BEAST2 (Table S2). This phylogeny resolved eastern oviparous, central viviparous, and western oviparous each as monophyletic (Fig. S2). However eastern viviparous, central viviparous, and western viviparous lineages were all polyphyletic, suggesting considerable introgression and a poor association of single gene mtDNA with the phylogeny generated from genome-wide data. Support values were generally considerably lower for both basal and terminal nodes compared to the phylogeny generated from the extensive genomic dataset. The topology also differed considerably from the topology generated from phylogenomic data (Fig. 3; Fig. S2).

**Figure 3.**
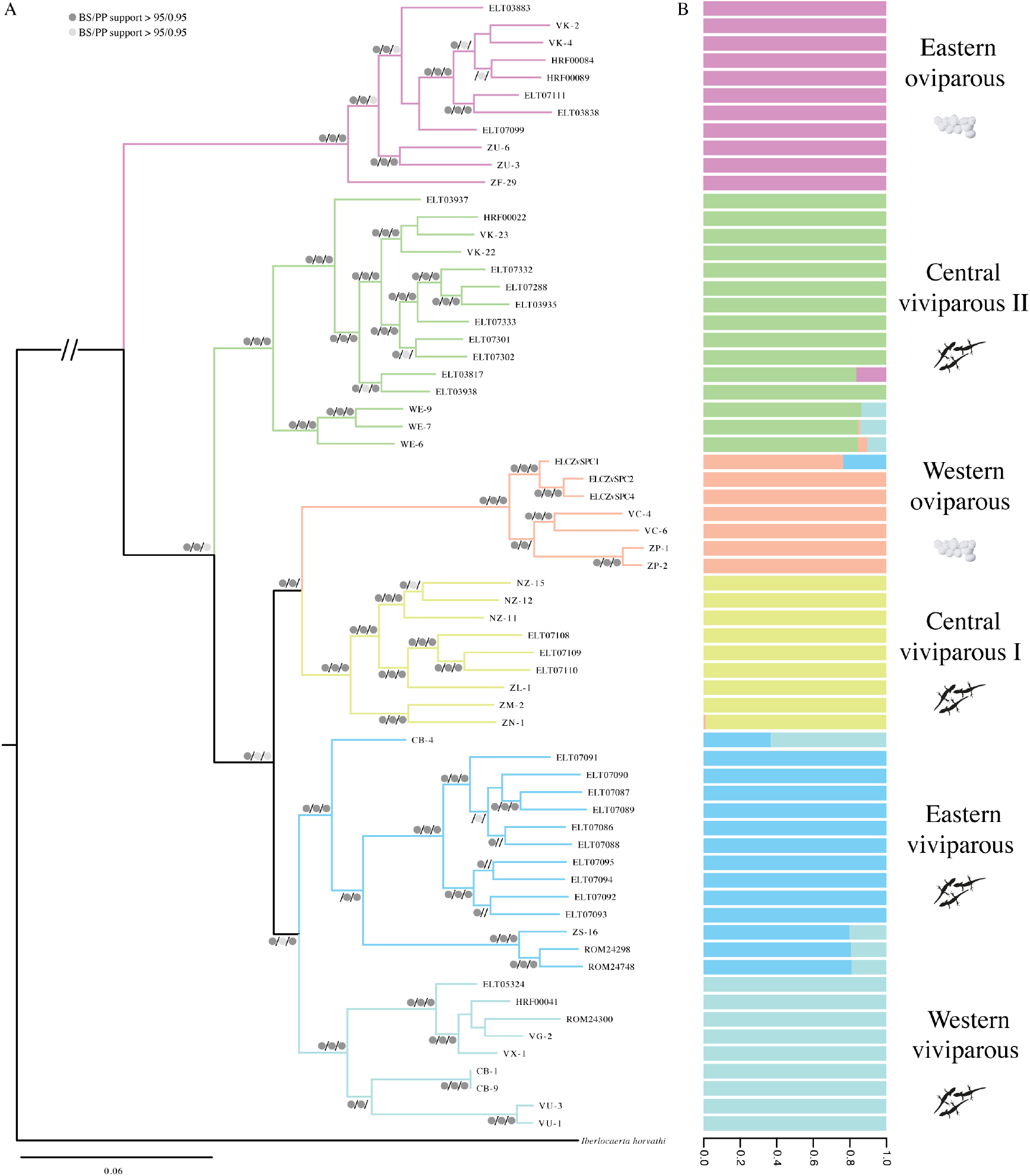
Bayesian (B), Maximum likelihood (ML) and maximum parsimony (MP) reconstruction of common lizard evolutionary relationships based on ddRADSeq data. A) The Bayesian tree was used with a full alignment using 1,334,760 sites (84,017 SNPs) and ML and MP trees were constructed with 194,358 SNPs. B posterior probabilities (BS), ML and MP bootstrap support are indicated by dark grey and light grey dots in that order (see legend). B) An ADMIXTURE analysis included the 194,358 SNPs and a k of 6 genetic clusters. Individuals are aligned vertically and respective membership values for each genetic cluster are illustrated. Parity mode and lineage are indicated on the right. *Iberolacerta horvathi* was used as an outgroup (true branch length not shown for graphical reasons).

### 3.3 *Monophyletic clades in* Zootoca vivipara *and reconstruction of evolutionary history*

All phylogenomic reconstructions confirmed six monophyletic evolutionary divergent lineages with high confidence (all MP and ML bootstrap supports of 100 and PP of 1.0; Fig. 3). The eastern oviparous lineage is basal sister to all other lineages, followed by central viviparous II. The remaining four lineages are split into two groups, one with the western oviparous and central viviparous I lineages as sister and one with the eastern and western viviparous lineages. This topology is concordant with a single origin of viviparity and a reversal to oviparity in the western oviparous lineage (see 3.2 for topological analyses). Population structure also confirmed these six genetic lineages, with high average membership values for each respective lineage (mean Q-values ranged from 92-100% identity within each lineages) (Fig. 3). These six lineages correspond to phylogeographic clades that were previously identified. The recently reported distinct Carpathian haploclade (Velekei et al., 2015) was not confirmed as a separate genetic cluster in our phylogenomic reconstruction and was nested within the Eastern viviparous lineage (individuals ELT07086-ELT07095). Our mitochondrial dataset confirmed monophyly of some of the lineages with good support (eastern oviparous, central viviparous, western oviparous), while others where not supported (Fig. S2). In contrast to the nuclear data, the separate Carpathian clade was strongly confirmed by mitochondrial DNA and monophyletic, sister to the eastern oviparous lineage (Fig. S2).

Genetic differentiation between all six lineages was substantial (Table S3). *Fst* and *Jost D’s* values were largest between eastern oviparous and all other lineages (Fst: 0.42 – 0.52; *Jost D*: 0.013 – 0.018), and second largest between western oviparous and all other lineages (*Fst*: 0.35 – 0.51; *Jost D*: 0.007 – 0.016), indicating that these are the most highly differentiated lineages. Compared to *Fst, Jost D* was weaker between the western oviparous and all other viviparous lineages (Table S3). Genetic differentiation between the viviparous lineages was less pronounced (*Fst*: 0.23 – 0.32; *Jost D*: 0.004 – 0.008).

### 3.4 Evolutionary scenarios for parity evolution

We found significant support for topologies associated with a single origin of viviparity and a reversal to oviparity. Bayesian, Maximum likelihood and parsimony analyses all confirmed the same topological configuration for the six main common lizard lineages with high nodal supports (bootstraps > 100, all posterior probabilities = 1.0) (Fig. 3). Phylogenies from all reconstruction methods support a topology in which the eastern oviparous lineage is basal to all other lineages. The following lineage splitting off is the central viviparous II lineage, sister to all remaining lineages. The western oviparous lineage is nested within the viviparous lineages, sister to the central viviparous I lineage. This topology suggests a single origin of viviparity in common lizards and a reversal to oviparity in the western oviparous lineage as the most parsimonious scenario for parity mode evolution.

Using monophyly constraints and statistical topology testing, any topologies compatible with alternative scenarios of parity mode evolution. Alternative scenarios included: oviparity as a basal trait and a single origin of viviparity (Figure 1A; Table 1), multiple independent origins of viviparity (Figure 1B; Table 1), a reversal to oviparity but independent sex chromosome evolution (Figure 1C; Table 1), and multiple origins of viviparity and a reversal to oviparity (Figure 1D; Table 1) and were all significantly less likely (Table 1) than a single origin of evolution, a reversal to oviparity and a single change in sex chromosome configuration, consistent with Figure 3.

**Table 1.** Statistics of alternative topological constraints. Five alternative topological constraints were set and compared to the best performing maximum likelihood tree. Topological constraints were set to represent different evolutionary hypotheses of parity mode evolution (assuming the most parsimonious path of evolution, i.e. the lowest number of possible transitions). Constraint models are ranked by observations, starting with the model without constraint. Constraint models are the following: i) ‘no constraint’ is consistent with a reversal to oviparity and refers to the topology in Figure 3, ii) ‘viviparous CVII basal’ is the same topology as i), only specifying the constraint that the central viviparous II lineage is sister to all remaining lineages excluding the eastern oviparous lineage, which is basal to central viviparous II; it is consistent with a reversal to oviparity and Figure 3, iii) ‘multiple viviparity’ constrains central viviparous II as sister to eastern oviparous, and western oviparous sister to all other viviparous lineages, consistent with two independent origins of viviparity and Figure 1B, iv) ‘oviparity basal’ constrains eastern and western oviparous lineages to be basal to all other viviparous lineages and is consistent with a single origin of viviparity and Figure 1A, v) ‘viviparous CVII not basal’ constraints the eastern oviparous lineage to be basal to all other lineages, but the central viviparous II not as basal to the remaining lineages; it is consistent with a reversal to oviparity but not with sex chromosome evolution and corresponds to Figure 1C, and vi) ‘viviparous RO basal’ constrains the Carpathian lineage to be sister to the eastern oviparous lineage, consistent with multiple independent origins of viviparity and potentially a reversal to oviparity and corresponds to Figure 1D.

| constraint | rank | obs | AU | NP | BP | PP | KH | SH | wtd-KH | wtd-SH |
| --- | --- | --- | --- | --- | --- | --- | --- | --- | --- | --- |
| no constraint | 1 | 0 | 0.518 | 0.493 | 0.502 | 0.500 | 0.496 | 0.918 | 0.496 | 0.918 |
| viviparous CVII basal | 2 | 0 | 0.535 | 0.501 | 0.494 | 0.500 | 0.504 | 0.891 | 0.504 | 0.891 |
| multiple viviparity | 3 | 404.6 | 0.000 | 0.000 | 0.000 | 0.000 | 0.000 | 0.021 | 0.000 | 0.000 |
| oviparity basal | 4 | 452.7 | 0.005 | 0.004 | 0.004 | 0.000 | 0.004 | 0.004 | 0.004 | 0.011 |
| viviparous CVII not basal | 5 | 1206.9 | 0.000 | 0.000 | 0.000 | 0.000 | 0.000 | 0.000 | 0.000 | 0.000 |
| viviparous ROM basal | 6 | 2478 | 0.000 | 0.000 | 0.000 | 0.000 | 0.000 | 0.000 | 0.000 | 0.000 |

Abbreviations are: obs = observations, AU = Approximately unbiased test, NP = non-scaled bootstrap probability, BP = bootstrap probability, PP = Bayesian posterior probability, KH = Kishino-Hasegawa test, SH = Shimodaira-Hasegawa test, wtd = weighted, CVII = central viviparous II, CVI = central viviparous I, RO = Carpathian viviparous clade.

Reconstructing evolutionary relationships between the six main phylogenetic lineages in TREEMIX results in a similar topology as retrieved from the other analyses, with eastern oviparous consistently sister to all other lineages. Overall likelihood and variance explained increased including more migration events, and reached a plateau after two migration events (Fig. S3). Topologies were unstable when more migration events were included, though these topological changes should be considered with caution since all *f*3-statistics were positive, indicating that admixture has not played a major role in the evolution of common lizard lineages (Table S4).

## 4. Discussion

### 4.1 Evolutionary history of parity mode evolution

Here, we show that the most parsimonious scenario for the evolution of parity mode evolution in common lizards includes a single origin of viviparity and a reversal to oviparity in a single lineage (western oviparous). Our genome-level phylogeny based on up to 194,358 nucleotides was highly supported by Bayesian ML, and MP analyses (support values >0.95). Topologies compatible with other parity mode scenarios, such as a no reversal to oviparity or multiple origins of viviparity (per Fig 1A, B, D) performed significantly worse in all statistical tests (Table 1). We find considerable differences between our high resolution phylogenomic tree and our mtDNA phylogeny.

The evolution of oviparity and viviparity in common lizards has been contentious and a range of studies, using different geographic and genetic sampling, have failed to converge on an evolutionary scenario. To date, mitochondrial DNA, nuclear DNA, and karyotypic markers have not agreed on a single topology (Fig. 1; Odierna et al., 2004; Surget-Groba et al., 2006, 2001; Velekei et al., 2015). For example, previous research suggested that a reversal to oviparity occurred in common lizards, however support was based on only limited data and support (Cornetti et al., 2014; Surget-Groba et al., 2006). It has also been proposed that viviparity evolved multiple times independently (Odierna et al., 2004; Velekei et al., 2015), however, these studies were limited to the use of a single marker. Our phylogeny is the first that is consistent with nuclear genetic markers and chromosomal configuration (Fig. 1; Fig. 3).

In addition to our robust and well supported phylogeny and the topological statistics, other aspects of common lizard genetics and reproductive traits also support our inference of a reversal to oviparity. The eastern oviparous and western oviparous lineages have different morphological and physiological egg characteristics, such as thinner eggshells and shorter incubation time (Arrayago et al., 1996; Lindtke et al., 2010). We suggest this is compatible with our phylogeny; the derived oviparous lineage is due to a reversal to oviparity instead of retaining the ancestral oviparous condition, and in doing so the thickness of the eggshell is reduced. Our phylogeny is consistent with the most parsimonious scenario for the derived chromosomal features in common lizards: While both the eastern oviparous and central viviparous II lineages have 36 chromosomes and a ZW sex chromosome configuration, all other lineages exhibit 35 chromosomes and a Z_1_Z_2_W sex chromosome configuration (Kupriyanova et al., 2008; Odierna et al., 2004; Fig. 1). Previous genetic studies were inconsistent with this derived sex chromosome configuration by placing central viviparous II nested within lineages exhibiting the Z_1_Z_2_W chromosome configuration instead of being basal to lineages with the derived configuration (Cornetti et al., 2014; Surget-Groba et al., 2001, 2006). The phylogeny presented here is the first molecular phylogeny consistent with a single transition in sex chromosome configuration, changing from the ancestral ZW system to the derived Z_1_Z_2_W system (Kupriyanova et al., 2006; Odierna et al., 2004).

Calcified eggshell and the associated reproductive life history traits of oviparity represent a complex character that once lost is unlikely to re-evolve, making it a trait long regarded to be subjected to Dollo’s law of irreversibility (Lee and Shine, 1998; Shine and Lee, 1999; Sites et al., 2011). However, research on the re-evolution of insect wings (Collin and Miglietta, 2008; Whiting et al., 2003), snail coiling (Collin and Cipriani, 2003), or mandibular teeth in frogs (Wiens, 2011) has shown that in some cases complex characters can indeed re-evolve. In squamate reptiles, one example exists arguing for the re-evolution of oviparity in sand boas (Lynch and Wagner, 2010). In this example, a scenario with no reversal to oviparity required three additional evolutionary transitions compared to the most parsimonious scenario with a single reversal to oviparity. In addition to the support from parsimonious trait reconstruction from the phylogeny, sand boas lack the egg tooth, which is an important anatomical structure for hatching from eggs that is present in related oviparous snake species. This provides independent evidence for the derived state in sand boas and the re-evolution of oviparity (Lynch and Wagner, 2010). In general, in addition to support from phylogenetic reconstruction, it should be best practice to assess whether the trait re-evolved is developmentally and anatomically similar to the ancestral trait. Substantially different features of the trait in the derived compared to ancestral form can be considered additional evidence for re-evolution, rather than the less plausible scenario that the ancestral form was retained but changed over time while an alternative trait was independently lost in multiple related lineages. In common lizards, the short timespan between the origin of viviparity and the reevolution of oviparity might have facilitated the reversal, in that not many genomic changes were required. In general, a trait as complex as viviparity is thought to require several changes in the genome (Murphy and Thompson, 2011).

Whether reversals to oviparity from viviparity occurred frequently in squamate reptiles remains a highly controversial topic. Erroneous phylogenetic reconstruction and limited assessment of characteristics of the trait in question have led to the publication of controversial examples of re-evolution (e.g. Fairbairn et al., 1998; Pyron and Burbrink, 2014) that have been criticized heavily (Blackburn, 1999, 2015; Griffith et al., 2015; King and Lee, 2015; Shine and Lee, 1999; Wright et al., 2015). Moreover, incomplete lineage sorting and/or introgression of the trait in question, combined with the limited molecular information included in most phylogenetic reconstructions, can lead to wrong conclusions in trait evolution (Hahn and Nakhleh, 2016). While here we found substantial support for the re-evolution of oviparity based on the largest genomic dataset to date, more knowledge on the development and genetics of the trait is necessary to unequivocally assess whether a reversal to oviparity occurred in common lizards. In the future, more refined phylogenetic reconstructions using whole genome and phylogenomic data combined with insights into the genetic mechanisms involved in parity mode evolution should provide answers on whether reversals to oviparity occur in squamates and how common they are.

### 4.2 Evolutionary relationships between common lizard lineages and comments on taxonomic status

Our genome-wide phylogeny recovered a new topology, but this included similar clades as previously supported by mitochondrial DNA reconstructions, except for the Carpathian clade, which we find is nested within the Eastern viviparous lineage (Fig. 1; Fig. 3; Fig. S3). Incongruence between nuclear data and mitochondrial data is observed frequently (Ballard and Whitlock, 2004; Near and Keck, 2013; Wallis et al., 2017). Consistent with previous phylogenetic analyses (Cornetti et al., 2014; Surget-Groba et al., 2006, 2001), we found the eastern oviparous lineage is basal to all other common lizard lineages. Splitting order for the other lineages differs from previous phylogenetic reconstructions, however, the reciprocal monophyly of all remaining five lineages was highly supported by all analyses here. In agreement with this, f3-statistics suggest that there was no significant admixture between lineages (Table S3). Past mitochondrial DNA introgression and capture are a possible mechanism explaining the discordance between mitochondrial and nuclear genes (Leavitt et al., 2017; Willis et al., 2014).

Based on the strong reciprocal monophyly of the lineages, we suggest that *Zootoca vivipara* should be divided into five or six subspecies. Some have argued that *Z v. carniolica* should be recognized as a separate species based on limited gene flow and reproductive isolation (Cornetti et al., 2015a, 2015b). However, while hybridization is rare and might be geographically restricted, it does occur between *Z v. carniolica* and other viviparous common lizards (Lindtke et al., 2010; pers. obs.) and phenotypic differences are generally small (Guillaume et al., 2006; Rodriguez-Prieto et al., 2017). Given the old evolutionary split (Surget-Groba et al., 2006) and its distinctive reproductive biology species status might be warranted. All other main lineages (CVII, CVI, EV, WV, WO) could each be rendered a subspecific status given their clear evolutionary splits and differences in karyotype (Guillaume et al., 2006; Kupriyanova et al., 2006; Odierna et al., 2004, 1998; Surget-Groba et al., 2006). Currently, only *Z. v. louislantzi* (WO) can be recognized as a valid subspecies, while other lineages have conflicting subspecific designations (Arribas, 2009; Schmidtler and Böhme, 2011). While diagnostic morphological features are scarce (Guillaume et al., 2006), in-depth analyses using more levels of the phenotype (e.g. differences in colouration, behavior, reproduction and ecology) should resolve whether the distinguished genetic lineages are supported by phenotypic data. A taxonomic revision for these lineages combined with morphological and ecological data across the whole distribution of the group is much-needed.

### 4.3 Advantages and challenges of RADSeq data for phylogenetic reconstruction

Our phylogenetic reconstruction represents the most comprehensive and robust phylogeny of common lizards to date, based on 194,358 bp of polymorphic SNPs and 67 individuals. Previous phylogenetic studies on common lizards using only mitochondrial data (Surget-Groba et al., 2006) or fewer nuclear markers (Cornetti et al., 2014) had only moderate congruency between different markers and weak support at basal nodes. In agreement with the challenges from previous studies, our mtDNA phylogeny of an established, informative locus was not compatible with the phylogenomic dataset, highlighting the limitations of mtDNA (Ballard and Whitlock, 2004; Wallis et al., 2017; Willis et al., 2014) and suggesting it is not an appropriate marker for resolving the history of common lizards. More generally, we suggest that for groups with short internal branches and evolutionary histories of recent to several million years divergence, the type of data produced by RADSeq might be optimal to resolve difficult evolutionary splits. This is the case for adaptive radiations or more generally for short and quick speciation events and complex phylogeographic histories(Giarla and Esselstyn, 2015; Rodríguez et al., 2017). This study evidences the power of fast evolving loci (loci with several SNPs) to resolve short phylogenetic branches.

A challenge of short-read phylogenomics and loci with multiple SNPs is the validity of orthology between loci. We show that topological groupings are more robustly supported when using loci with multiple SNPs (Fig S1) and we present an assessment pipeline for validating the cut-offs for missing data and SNPs per locus. Without a reference genome and a large amount of duplicated and/or repetitive DNA, orthology of RAD loci is usually not evaluated. Using a reference genome to map the RAD loci and high sequencing coverage per individual, such as done here, are important methodological considerations to overcome these issues (Mastretta-Yanes et al., 2015; Shafer et al., 2017). Disadvantages of these large but informative datasets are long computational time for some analyses, in particular phylogenetic reconstructions using Bayesian coalescence based analyses (Bryant et al., 2012). Advances in phylogenomic methodologies to accommodate these more complex datasets will be important for advancing the field (Delsuc et al., 2005; Fuentes-Pardo and Ruzzante, 2017; Leavitt et al., 2016).

### 4.4 Conclusions

Our results strongly support a single origin of viviparity in common lizards and a subsequent reversal to oviparity in one derived lineage as the most parsimonious scenario of reproductive mode evolution (Fig 3, Table 1). In the light of karyological and reproductive data (Arrayago et al., 1996; Heulin et al., 2002; Lindtke et al., 2010; Odierna et al., 2004, 1998), these findings are strong evidence that a reversal to oviparity has occurred what is now the allopatric western oviparous lineage (Fig. 2, Fig. 3). In addition, we propose that a taxonomic revision of this genus at the subspecific level may be needed. More generally, this suggests that Dollo’s law of irreversibility is not without exceptions, and might be particularly prone to switches between characters at early stages of evolution of a new or lost trait. For the future, we suggest that common lizards represent an ideal candidate to investigate the genomic basis for evolutionary complex reversals.

## Acknowledgments

This work would not have been possible without the support and contribution of samples by Werner Mayer, to whom we are very grateful. We thank B. Murphy and A. Lathrop at the Royal Ontario Museum for providing tissue samples. We particularly thank Austrian and Scottish authorities for issuing collection permits (HE3-NS-959/2013; SNH license number 64972). We thank Megan Layton, Henrique Leitão, Mark Sutherland, Ruth Carey, Michael Andrews, Jade McClelland, and Nathalie Feiner for assistance and companionship in the field during the collection of crucial samples. We thank Aileen Adams, Arne Jacobs, Julie Galbraith, Jing Wang, Lorraine Glennie, and Peter Jeffrey Koene for their help in the lab and A. Yurchenko for access to the reference genome and valuable discussions. For funding we gratefully acknowledge a Heredity Fieldwork grant by the Genetic Society to HR, a University of Glasgow Lord Kelvin-Adam Smith PhD Studentship to KRE and NK for HR, and NERC grant NE/N003942/1 to KRE.

## Author Contributions

KRE, NK and HR conceived the study. HR and KRE collected samples and designed the experiments. HR generated data, performed all analyses and drafted the manuscript. KRE, NK and HR all contributed to the writing of the final version of manuscript.

## Conflicts of Interest

The authors declare no conflict of interest.

